# Temporal-Spatial Visualization of Endogenous Chromosome Rearrangements in Living Cells

**DOI:** 10.1101/734483

**Authors:** Haifeng Wang, Muneaki Nakamura, Dehua Zhao, Cindy M Nguyen, Cordelia Yu, Albert Lo, Timothy Daley, Marie La Russa, Yanxia Liu, Lei S Qi

**Affiliations:** Department of Bioengineering, Stanford University, Stanford, CA 94305, USA; Castilleja School, Palo Alto, CA 94301, USA; Department of Statistics, Stanford University, Stanford, CA 94305, USA; Department of Chemical and Systems Biology, Stanford University, Stanford, CA 94305, USA; ChEM-H Institute, Stanford University, Stanford, CA 94305, USA

## Abstract

Visualizing the real-time dynamics of genome rearrangement in single living cells is core to studying genomics and diagnostics. Here, we report a robust, versatile approach named CRISPR <u>Live</u>-cell fluorescent *in <u>s</u>itu* <u>h</u>ybridization (LiveFISH) for multi-locus genome tracking and cytogenetic detection in a broad variety of cell types including primary cells. LiveFISH utilizes an intrinsic stability switch of CRISPR guide RNAs, which enables efficient and accurate detection of chromosomal disorders such as Patau Syndrome in prenatal amniotic fluid cells and allows multi-locus tracking in human T lymphocytes. Using LiveFISH, we are able to detect and track real-time spatiotemporal dynamics of non-homologous endogenous chromosome translocations induced by gene editing. This new approach enables FISH imaging in living primary cells, which can provide useful insights into the spatiotemporal changes of genome organization and rearrangements in normal and diseased primary cells and will enable fast cytogenetic visualization of various gene-editing associated chromosomal translocations.

## Introduction

Large-scale chromosomal rearrangements occur in various genetic disorders and cancers (1). For example, chromosome duplications cause Down syndrome and Patau Syndrome (2,3), and many cancers including chronic myelogenous leukemia and Burkitt’s lymphoma are correlated with large-scale chromosomal translocations (4,5). Genome editing in human cells was recently reported to induce long-range rearrangement events including inversions and translocations (6,7). Despite efforts to identify and characterize chromosomal abnormalities associated with genetic disorders or with deliberate manipulations of the genome such as gene editing, little is known about spatiotemporal dynamic process leading to endogenous chromosomal rearrangements. Traditional cytogenetic diagnosis has relied on chromosomal karyotyping and fluorescent *in situ* hybridization (FISH) (8), which are restricted to non-living cells thereby precluding the ability for spatiotemporal analyses in living cells. Here, we aim to develop a live-cell genomic imaging approach that allows multi-locus detection and real-time study of endogenous chromosomal rearrangements.

Visualizing genomic rearrangements in living cells using pre-integrated LacO and TetO arrays provides valuable mechanistic information (9). However, this method is unsuitable for studying endogenous translocations and making the required cell lines is time-consuming (9). The use of the nuclease-deactivated dCas9 (CRISPR-associated protein 9) derived from type II CRISPR (clustered regularly interspaced palindromic repeats) system coupling with fluorescent components has allowed more flexible genome imaging in living cells (10–20). However, creation and optimization of cell lines stably expressing all CRISPR components in each cell type are often required to achieve robust imaging, which is challenging in primary cells (21,22). CASFISH has been developed to use in vitro assembled CRISPR complexes to detect genomic loci, which however, still requires fixation and permeabilization of cells and tissues (23). New approaches are needed for the cytogenetic detection of disease- or gene-editing-associated chromosomal abnormalities in living primary cells, and for the in-depth analysis of mechanisms underlying the dynamic process of chromosomal rearrangements.

Here, we present an approach we call CRISPR LiveFISH, for robust real-time visualization of endogenous chromosomal rearrangements such as duplications and translocations in living primary cells. Taking advantage of an intrinsic stability switch of CRISPR guide RNA (gRNA), we used fluorescent ribonucleoproteins (fRNPs) consisting of fluorescently labeled gRNAs and the dCas9 protein for imaging in living cells (24,25). We show that LiveFISH allows fast and robust cytogenetic detection of Patau Syndrome in patient amniotic fluid cells, multiple chromosomal loci tracking in primary T lymphocytes, and real-time visualization of chromosome translocations in single living cells induced by CRISPR gene editing.

## Results

### Target DNA binding stabilizes gRNAs in the Cas9:gRNA:DNA ternary complex in the RNase-rich environment *in vitro* and in cells

The Cas9 protein is directed to the DNA target by a gRNA duplex composed of a CRISPR RNA (crRNA) and a trans-activating crRNA (tracrRNA), or a single gRNA (sgRNA) in which the two RNAs are fused together (11). The gRNA has shown to be highly unstable in cellular environments (26,27). For example, gRNA is extremely unstable without Cas9 binding (undetectable in cells), and it still exhibits a short half-life (15 min to 2 hours) even with Cas9 binding compared to that of the Cas9 protein (’12 hours) (26–28). While efficient Cas9-mediated gene editing and regulation require increased stability of gRNAs (16,26,27), we reasoned that imaging applications can utilize their unstable nature. We first investigated how gRNA stability is affected by gRNA interaction with Cas9 and on-target or off-target DNAs in the RNase-rich environment *in vivo* and *in vitro*.

To do this, we assembled equimolar amounts of dCas9-EGFP protein and Cy3-labeled gRNA duplex consisting of both Cy3-crRNA and tracrRNA into fRNPs, which target a repetitive region within Chromosome 3 (Chr3) (11,16,18). We delivered the fRNPs into human bone osteosarcoma U2OS cells via electroporation (**Figure 1A**) (29). We observed rapid recruitment of Cy3-gRNA to the Chr3 target site after transfection, which remained detectable after 72 hours. This suggests formation of stable DNA target-bound dCas9-gRNA complexes (**Figure 1B**). Unlike dCas9-EGFP that frequently accumulated in the nucleolus, there was no nucleolar accumulation of gRNA (**Figure 1C**). Time-course flow cytometry analysis confirmed that more than 95% of gRNA signals were degraded within 4 hours after transfection (**Supplemental Figure 1A**), which was consistent with previous reports on the instability of gRNAs in cells (26,27). The ability to detect on-target gRNA signals for 72 hours implies that dCas9-bound gRNA is highly stable when bound to its cognate DNA targets.

**Figure 1.**
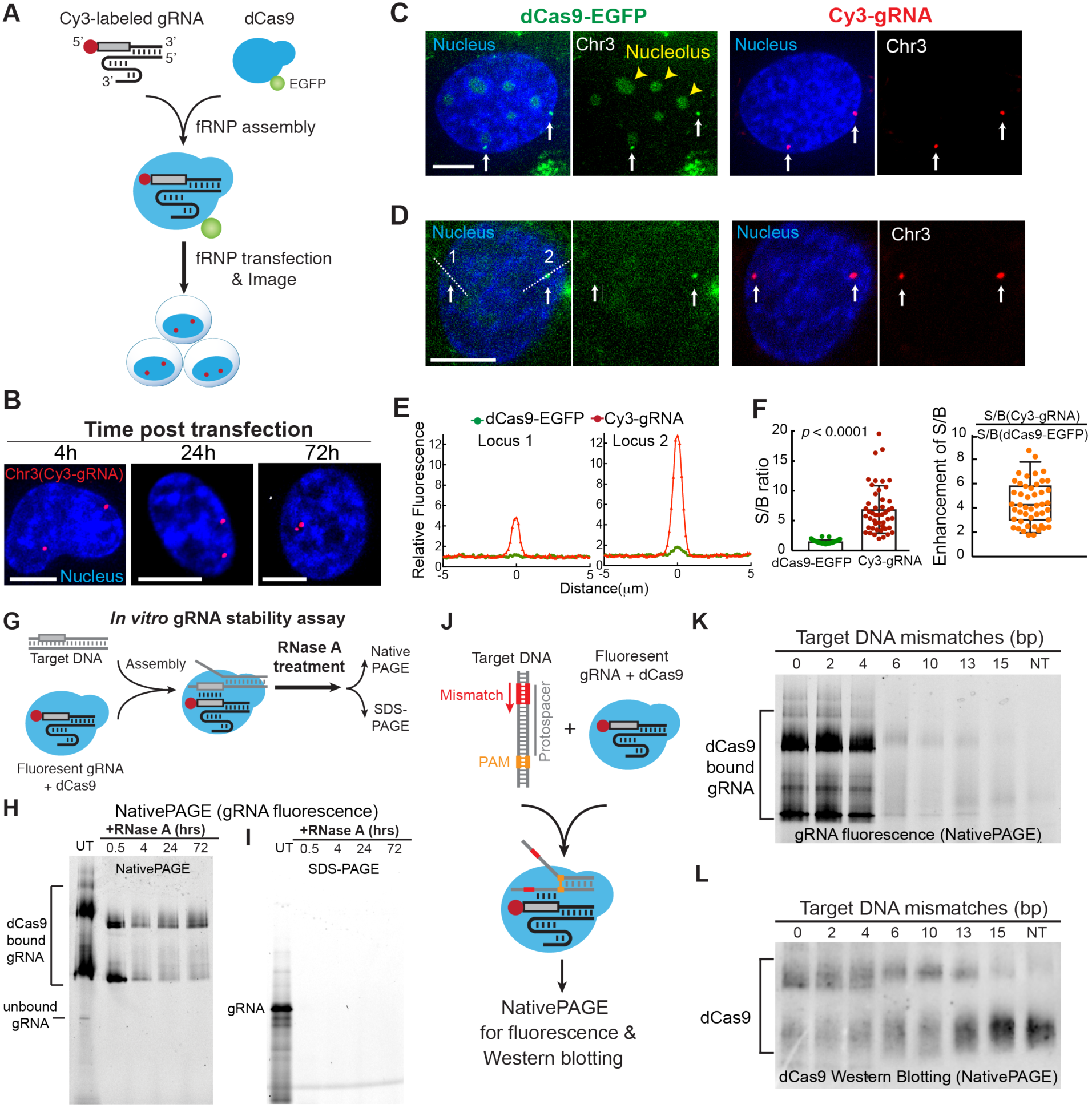
Target DNA binding stabilizes gRNAs in vitro and in cells. (**A**) Schematic of a fluorescent ribonucleoprotein (fRNP) approach to study the stability of fluorescent gRNAs in living cells. Fluorescent gRNAs (cr:tracrRNAs) were assembled with dCas9 or dCas9-EGFP proteins as fRNPs and then transfected into cells. Cy3 labels the 5’-end of crRNA. (**B**). Fluorescent microscopic images showing representative U2OS cells with Cy3-gRNA targeting an endogenous chromosome 3 (Chr3) loci at different time points (4, 24 and 72 hours) after electroporation. Hoechst 33342 (blue) labels the cell nucleus. Scale bars, 10 μm. (**C, D**) Comparison of fluorescent gRNA (Cy3-gRNA, red) and dCas9-EGFP (green) labeling for Chr3 (white arrows) in U2OS cells via fRNPs. Hoechst 33342 (blue) labels the nucleus. The yellow arrowheads in **C** show dCas9-EGFP accumulation in nucleolus. The white dotted lines in **D** are used for linescan data in **E**. Scale bars, 10 μm. (**E**) Linescans of the cell shown in **D** (white dotted lines) to compare signal-to-background (S/B) ratio for each Chr3 locus in Cy3-crRNA (red) and dCas9-EGFP (green) channels. Relative fluorescence is calculated by setting average background fluorescence as 1. (**F**) Comparison of S/B ratio of labeled Chr3 loci using fluorescent gRNA (red) and dCas9-EGFP (green). Left: data are represented as mean ± SD and P<0.0001 by *t*-test. Right: the calculated ratio (orange) between the S/B of Cy3-crRNA and dCas9-EGFP channels at each locus. Average, SD, 5% and 95% percentiles are shown in right. 47 loci in 17 cells are analyzed. (**G**). Schematic of a gRNA stability assay testing the stability of fluorescent gRNAs in complex with dCas9 and on-target DNAs. Fluorescent gRNAs in complex with dCas9 were incubated with on-target DNAs, and then treated with RNase A. Samples were taken after RNase A treatment and run on native (NativePAGE) and denaturing (SDS-PAGE) electrophoresis. (**H-I**) Native (**H**, NativePAGE) and denaturing (**I**, SDS-PAGE) gel electrophoresis analyses compare the fluorescence of Atto565-gRNAs in complex with dCas9 and on-target DNA treated with RNase A for 0.5, 4, 24, and 72 hours (right four lanes). UT, Untreated with RNase. Atto565 labels 5’-end of crRNA. Fluorescent gRNAs were detected by fluorescence. (**J**) Schematic of a biochemical assay testing how target DNA mismatches affect gRNAs in complex with dCas9. Fluorescent gRNAs were assembled with dCas9 and incubated with on-target DNA or DNAs containing mismatches at the PAM-distal end. Samples were analyzed by native electrophoresis (NativePAGE). Fluorescent gRNAs were detected by fluorescence, and dCas9 proteins were detected by Western blotting. (**K-L**) NativePAGE analysis of fluorescent Atto565-gRNAs (**K**, gRNA fluorescence) in complex with dCas9 (**L**, Western blotting) incubated with different DNA duplexes. Lanes labeled with “0”, “2”, “4”, “6”, “13”, “15” and “NT” show Atto565-gRNA in complex with dCas9 incubated with DNA comprising 0, 2, 4, 6, 10, 13, or 15bp PAM-distal mismatches or a non-targeting DNA.

The gRNA signal showed a much higher signal-to-background ratio (S/B, by 4.4±1.8 fold) compared to the dCas9-EGFP signal (**Figure 1D-F, Supplementary Figure 1B)**. The S/B ratio is largely determined by the concentrations of local DNA target-bound fluorophores versus global unbound fluorophores. The improved S/B ratio using Cy3-gRNA can be explained by selective protection of the on-target gRNA and rapid degradation of unbound gRNA, as well as other factors including advantages of using chemical fluorescent dyes over fluorescent proteins and the differences in cell autofluorescence.

To confirm the DNA-dependent protection of gRNA, we performed *in vitro* assays to test whether on-target DNA binding can stabilize the gRNA within the Cas9:gRNA:DNA ternary complex against RNase degradation. fRNPs containing dCas9 and Atto565-labeled gRNA were incubated with excessive amounts of targeting or non-targeting DNA and RNase A, and analyzed by the native gel electrophoresis (NativePAGE) (**Figure 1G-H**). In the presence of the targeting DNA, dCas9-bound fluorescent gRNAs remained stable after 72 hours of incubation at 37°C (**Figure 1H**). In contrast, using the non-targeting DNA we observed rapid gRNA destabilization (**Supplementary Figure 2A**). To confirm that protection of gRNA within the Cas9:gRNA:DNA ternary complex relied upon binding to dCas9, we ran the same samples from **Figure 1H** with SDS-PAGE, an assay that disrupted dCas9 structure (**Figure 1I**). No fluorescent gRNA was detected in RNase A-treated samples, suggesting rapid degradation of gRNA by residual RNase A after dCas9 denaturation during sample preparation and electrophoresis. These results together suggest a strong protection of gRNA against RNase degradation when complexed with both dCas9 and the on-target DNA molecule.

To investigate how DNA mismatch affects gRNA stability, we incubated the fRNP for 30 minutes with DNA targets containing a variable number of mismatches, and analyzed the samples on NativePAGE (**Figure 1J-L)**. While similar total amounts of dCas9 were detected by Western Blot in each sample (**Figure 1L**), dCas9-bound gRNA exhibited high fluorescence with the fully matched target DNA or target DNAs with ≤4nt PAM-distal mismatches, but low fluorescence in the presence of target DNAs with ≥6nt mismatches (**Figure 1K)**. We propose that target DNA binding with ≤4nt mismatches stabilizes the crRNA within the Cas9:gRNA:DNA ternary complex, and that this protection is lost with DNA containing ≥6nt mismatches. As a control, we found that the total fluorescence in solution exhibited no difference in samples containing the targeting DNA or the non-targeting DNA (**Supplementary Figure 2B**), suggesting that target DNA binding does not directly alter the brightness of fRNPs.

These results suggest that the gRNA is protected in the Cas9:gRNA:DNA ternary complex under the RNase-rich environment *in vitro* and in cells. The protection mechanism is probably associated with conformational changes of Cas9 upon full gRNA and DNA binding (30–36), and is consistent with studies showing that dCas9 stays bound to on-target genomic loci much longer than to off-target loci in cells (27,37).

### Target DNA-dependent gRNA stabilization enables CRISPR LiveFISH, a rapid, accurate, and multiplexed approach for genomic visualization and diagnosis in living primary cells

Based on this target DNA-dependent protection mechanism, we developed CRISPR LiveFISH (live cell fluorescent *in <u>s</u>itu* <u>h</u>ybridization), an approach that deploys fluorescent gRNA probes assembled with dCas9 to hybridize with targeted genomic sequences in living cells for genomic visualization and cytogenetic diagnosis (**Figure 2A**).

**Figure 2.**
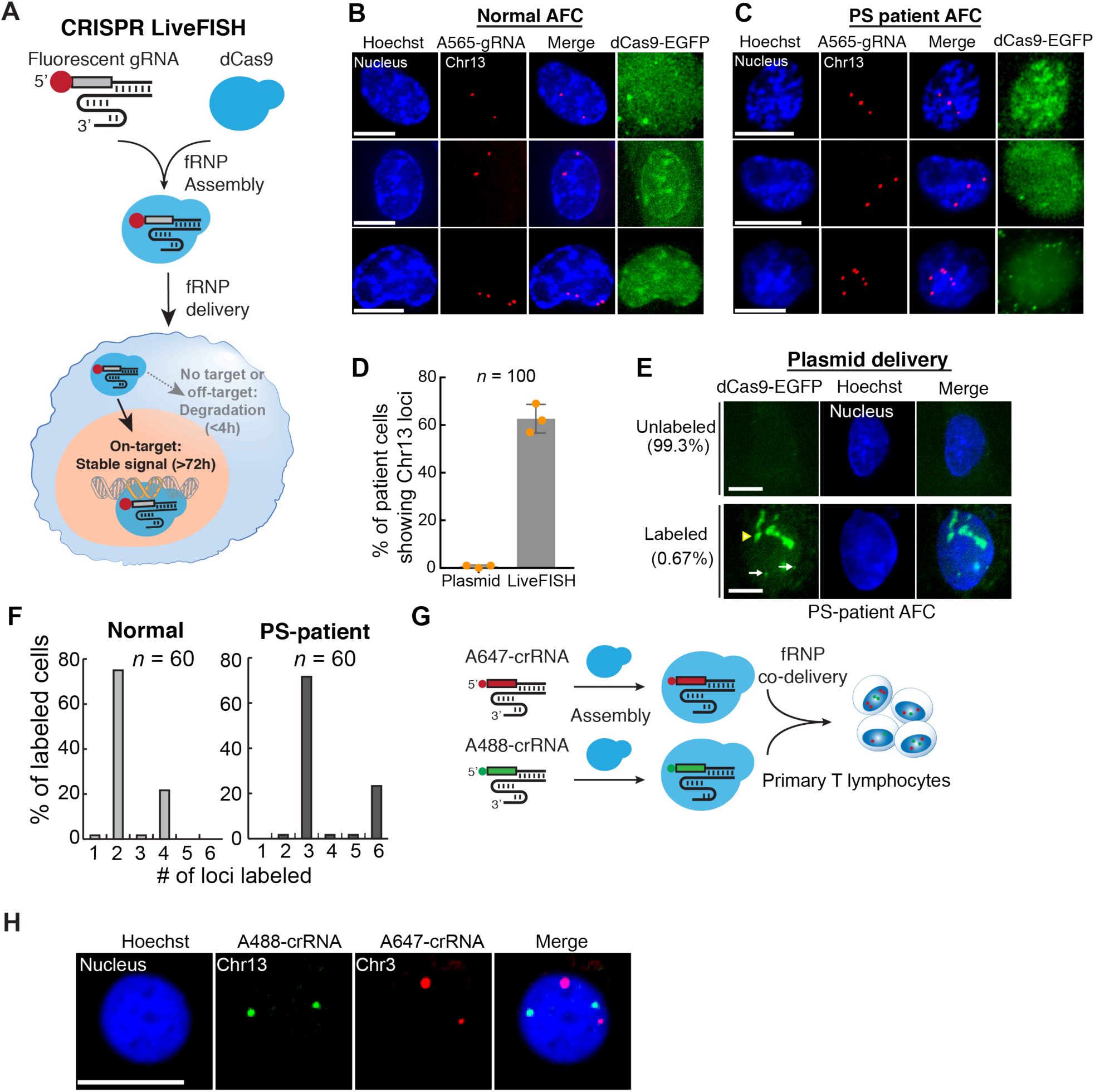
DNA target-dependent protection of gRNA enables CRISPR LiveFISH for genomic tracking and cytogenetic diagnosis in primary cells. (**A**) Schematic of the CRISPR LiveFISH approach for genomic imaging and cytogenetic detection in living cells via fRNP consisting of synthesized fluorescent gRNAs and dCas9. The dCas9 protein mediates hybridization between fluorescent gRNA probes and targeted genomic DNAs. The fluorescent gRNA is only stabilized upon binding to the genomic targets, and unbound gRNAs are rapidly degraded to reduce the background. (**B, C**) Images of CRISPR-LiveFISH genomic imaging and cytogenetic detection in normal (**B**) and PS patient-derived (**C**) amniotic fluid cells (AFCs) with Atto565-gRNA targeting Chr13 (red). Hoechst 33342 (blue) labels the nucleus. Cells were transfected with fRNPs containing dCas9-EGFP (green) and Atto565-gRNA (red). See **Movies S1 & S2** for dynamics. (**D**) Comparison of the percentage of PS-patient AFCs showing labeled Chr13 loci using DNA-encoded dCas9-EGFP and LiveFISH. Data are represented as mean ± SD. ≥ 100 cells from the same patient were analyzed for each of three independent experiments. *p*=0.0027 by a *t-*test. (**E**) PS patient-derived AFCs transfected with plasmids coding TRE3G-dCas9-EGFP and sgRNA targeting Chr13. Only 0.67% of cells showed labeled Chr13 loci (white arrows). Nucleolar accumulation is indicated by a yellow arrowhead. Hoechst 33342 labels the nucleus (blue). (**F**) Histograms of the number of Chr13 loci detected in normal and PS patient-derived amniotic fluid cells by LiveFISH. 60 cells from each group were analyzed. (**G**) Schematic of LiveFISH for multi-locus genomic imaging using multiple fRNPs. crRNAs targeting different genomic loci are synthesized with different fluorophores, annealed with tracrRNAs, and assembled with dCas9 to form multi-color fRNPs. (**H**) Simultaneous labeling of Chr13 (Atto488-crRNA, green) and Chr3 (Atto647-crRNA, red) by co-delivery of two fRNP complexes in living human T lymphocytes. Hoechst 33342 (blue) labels the nucleus. See another example in **Movie S3** for dynamics. Scale bars, 10 μm.

We tested whether LiveFISH can be used to detect chromosome abnormalities and track genomic loci in primary cells. We chose Patau Syndrome (PS) as a test case, which is a genetic disorder caused by three copies of Chromosome 13 (Chr13) that results in organ defects, intellectual and physical impairment, and is eventually fatal to patients (2). We transfected fRNPs containing dCas9 and Atto565-labeled gRNA to target Chr13 into normal or PS patient-derived prenatal amniotic fluid cells (AFCs). After four hours of transfection, we detected rapid and robust Chr13 labeling in the Atto565-gRNA channel in living normal (**Figure 2B, Movie S1**) and PS patient-derived AFCs (**Figure 2C, Movie S2**).

We compared LiveFISH with RNP and DNA delivery of dCas9-EGFP for genomic imaging in primary cells. While Atto565-gRNA robustly visualized Chr13 loci in 62% of AFCs (n=100 cells in 3 duplicates, **Figure 2D**), both RNP (**Figure 2B-C**) and DNA (**Figure 2E**) delivery of dCas9-EGFP showed almost no Chr13 labeling (<1%) in AFCs due to high background and nucleolar aggregation. Thus, LiveFISH is advantageous to previous CRISPR imaging methods for genomic imaging in primary cells. This also confirmed that the robustness of LiveFISH in primary cells is due to the selective protection of on-target gRNAs: gRNAs are stabilized upon binding to the genomic target, which preserves signals over a long period of time. In contrast, unbound or off-target bound gRNAs are rapidly degraded to eliminate background signal (**Figure 2A**).

We quantified the accuracy and efficiency of LiveFISH for cytogenetic imaging in living primary cells. Among labeled cells 97% of normal AFCs showed two (75%) or four (22%) Chr13 loci, and 95% of PS patient-derived AFCs showed three (72%) or six (23%) loci (**Figure 2F**). These numbers were consistent with expected copy number of chromosomes in each cell population at different phases of the cell cycle (G0/G1 or S/G2/M), which was confirmed by karyotyping (**Figure 3A-B, Supplementary Figure 2C-D**).

**Figure 3.**
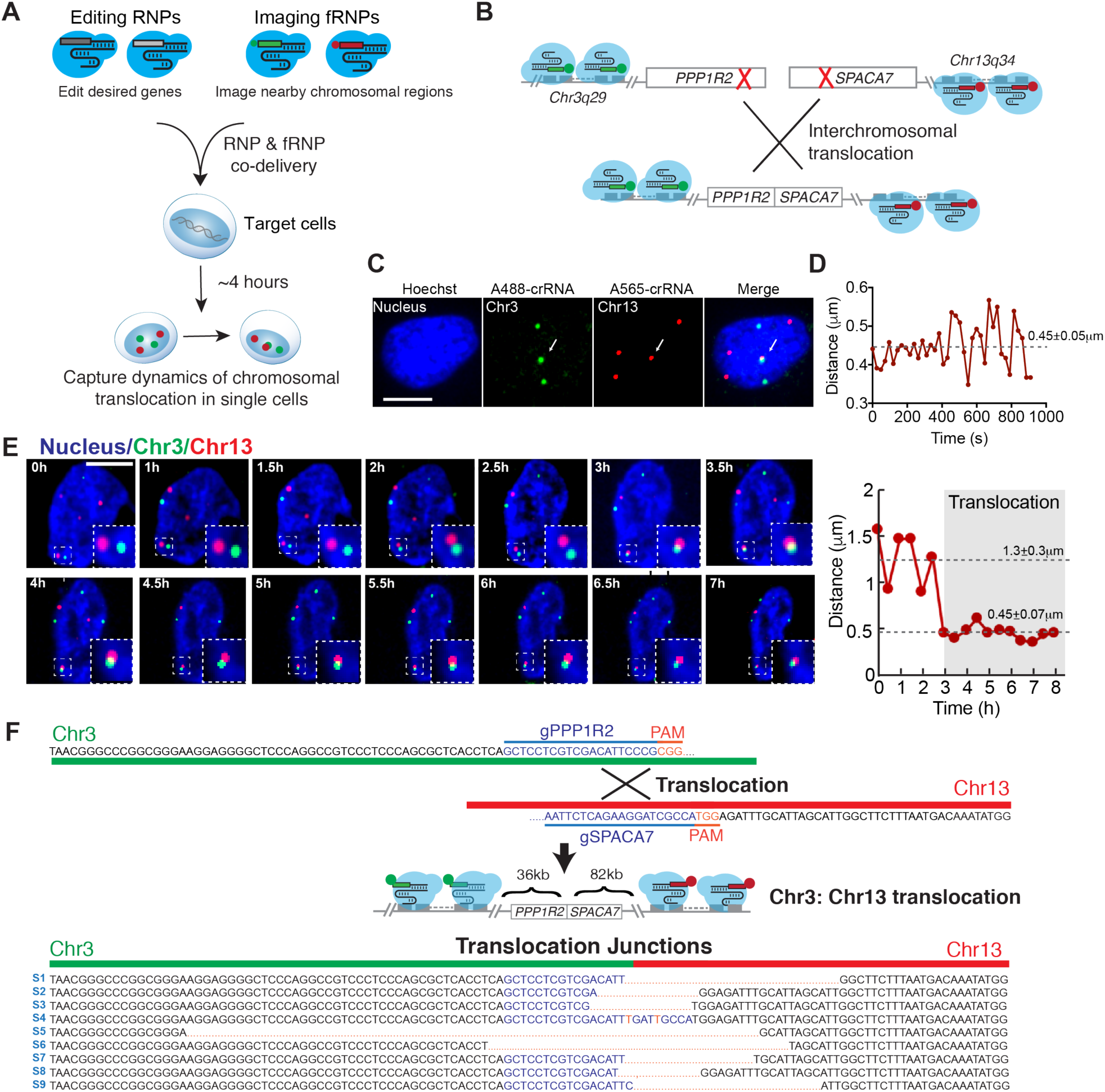
Real-time detection and visualization of non-homologous chromosome translocation in living cells. (**A-B**) Schematic of visualization of chromosomal translocation by combining LiveFISH fRNP complexes with CRISPR gene editing RNP complexes. Cas9 RNPs were assembled with gRNAs cutting *PPP1R2* on Chr3 and *SPACA7* on Chr13, and two imaging gRNAs targeting chromosomal regions 18-82kb away from the edited sites (Chr3: Atto488-crRNA, green; Chr13: Atto565-crRNA, red). The cutting sites are shown in red crossings in **B**. (**C**). A representative U2OS cell showing Chr3 and Chr13 loci paired together (arrow), likely representing a translocated chromosome. (**D**) Distances between the paired Chr3 and Chr13 loci over time shown in C. (**E**) A representative U2OS cell showing the formation process of chromosomal translocation between Chr13 and Chr3 loci (outlined in white boxes). (**F**) Sequences of amplified Chr3:Chr13 translocations around cutting gRNA targeting sites. gRNA targeting sites are shown in blue and PAMs are shown in red. 9 different translocated segments (S1-S9) were detected in 24 sequenced clones. Mismatches (red) show missing or altered nucleotides within the translocated junctions likely due to resections. See **Supplementary Figure 5D** for raw sanger sequencing data. Scale bars, 10 μm.

LiveFISH also simplifies visualization of multiple genomic loci in primary cells. We mixed individual fRNPs containing Atto647-crRNA targeting Chr3 and Atto488-crRNA targeting Chr13 and co-delivered them into primary human T lymphocytes (**Figure 2H, Supplementary Figure 4**). After 6 hours, we detected Chr13 and Chr3 labeled with different colors in the same cells (**Figure 2I**). We were able to track the dynamics of each genomic locus through time-lapse microscopic imaging (**Movie S3**). Compared to previous methods using multiple orthogonal dCas9s (17,22) or coupling sgRNAs to RNA aptamers and RNA binding proteins (18,38), which necessitated usage and optimization of many constructs, LiveFISH greatly facilitates multi-locus visualization via simple co-delivery of multiple fRNPs.

### Real-time visualization of gene editing-induced non-homologous chromosome translocations in living cells

We next applied the real-time multi-locus tracking ability of LiveFISH to capture chromosome translocation induced by gene editing. The use of CRISPR to edit multiple genes has important implications for gene therapy and cell therapy (39,40), but can induce non-homologous chromosomal rearrangements that may have long-term adverse effects such as oncogenesis (6,7). To our knowledge, no approach is available to capture and visualize the process of endogenous chromosomal translocation in individual living cells.

To visualize chromosomal translocations induced by CRISPR gene editing, we used Cas9-gRNA RNPs to edit two genes, *PPP1R2* on Chr3 and *SPACA7* on Chr13, and simultaneously co-delivered LiveFISH fRNPs to visualize the dynamics of both chromosomes (**Figure 3A-B**). The fRNPs contained two fluorescent gRNAs (Atto488-crRNA for *PPP1R2* and Atto565-crRNA for *SPACA7*) that labeled chromosomal regions 18-82kb away from the edited sites (**Figure 3B**). This one-step approach allowed us to simultaneously edit genes and track editing-associated chromosomal translocations in individual living cells. To avoid unwanted fRNP-mediated DNA cutting, the imaging gRNAs possessed short (11nt) spacers, which were insufficient for DNA cleavage (18,41).

We first confirmed double-stranded break (DSB) formation at targeted genomic loci by observing co-localization of fRNP signals with the DNA damage marker, 53BP1, 10 hours after transfection (**Supplementary Figure 5A**). As reported before, we found U2OS cells presented extreme aneuploidy (42). Using CRISPR LiveFISH, we tracked the short-term (15 minutes) dynamics of paired Chr3 and Chr13 loci (’6% of U2OS cells, n=363) and observed almost identical movement trajectories (**Movie S4**). The distance between the paired chromosomal loci remained almost constant (0.45±0.05µm) with small variations, which suggests a stable physical link between the two chromosomal loci (**Figure 3C-D**).

To capture the real-time dynamics of chromosomal translocation, we tracked the labeled loci over a long period of time in individual cells. Shown as an example in **Figure 3E and Movie S5**, a Chr3 locus and a Chr13 locus were originally spaced ’1.5µm apart at t = 0, then subsequently moved closer at t = 3h, and stayed together for the remaining observation period (over 5 hours). The two loci were spaced 1.27±0.29µm and 0.45±0.07µm before and after t = 3h, respectively, which likely represents an event of the endogenous chromosomal translocation formation (**Figure 3E & Movie S5**).

To independently confirm chromosomal translocation, we cloned and sequenced the translocated segments according to reported PCR-based methods (6,7). Multiple pairs of PCR primers confirmed translocated chromosomal segments with expected lengths in gene edited samples, while no translocated segments were detected in unedited cells (**Supplementary Figure 5B-C**). These sequencing results confirmed a number of translocation events between endogenous Chr3 and Chr13, which generated non-homologous chromosomal fusions containing LiveFISH-labeled Chr3 and Chr13 segments that are ’118kb apart with indels at the translocation junctions (**Fig. 3F, Supplementary Figure 5D**).

## Discussion

Here, we report a robust and versatile technology for live-cell imaging, CRISPR LiveFISH, which can visualize chromosomal dynamics in a broad variety of cell types including primary cells. LiveFISH utilizes chemically synthesized fluorescent gRNAs in fRNP complexes to facilitates rapid (a few hours), robust (a high S/B ratio), and multi-locus genomic tracking in living cells, which combines the advantages of FISH (easy-to-use, high sensitivity, multiplexibility, versatility in cell types) and CRISPR imaging (unfixed samples, live-cell imaging) (**Supplementary Figure 6)**. As a versatile method, a wide range of probes (e.g., quantum dots, biotin, and heavy metals) can be conjugated to gRNAs for various applications.

We characterized the target DNA-dependent protection of gRNAs within the Cas9:gRNA:DNA ternary complex in the RNase-rich environment. This mechanism enables enrichment of on-target gRNA signals while minimizing off-target or background signals. We applied LiveFISH for real-time genomic tracking and cytogenetic detection of chromosomal duplications in Patau Syndrome patient-derived cells. We showed that, while previous CRISPR imaging methods failed to work, LiveFISH enabled rapid, accurate, and multiplexable chromosomal abnormality detection in living primary samples, which will facilitate fundamental genomic research and point-of-care diagnosis of genetic orders and cancers.

We further used LiveFISH for real-time visualization of gene editing-induced chromosomal translocation events. To our knowledge, this represents the first demonstration of tracking the dynamics of CRISPR-induced translocation between endogenous chromosomes in living cells. Chromosomal translocations or rearrangements present a major concern for gene editing applications in therapeutics, and they are often associated with a variety of diseases and cancer. Detecting low-frequency chromosomal translocations and their formation process is challenging using existing methods (e.g., FISH), which are either restricted to non-living samples, or require a process (e.g., LacO/TetO arrays) to generate stable cell lines that makes it unsuitable for studying endogenous translocations in primary samples (9). LiveFISH provides a unique solution for real-time visualization of endogenous chromosome rearrangements via a one-step reaction. For example, the fRNPs can be transiently introduced into cells with gene editing molecules, which will be subsequently degraded with no long-term adverse effects on the imaged samples. Cells can be recovered after LiveFISH analysis for culture, sequencing, and functional characterization (**Supplementary Figure 7**). The human genome contains many repetitive sequence clusters that are highly abundant and adjacent (within 100kb) to the majority (>60%) of encoding genes, which can serve as robust chromosomal imaging regions (43). Given the targeting-flexibility of LiveFISH, our approach offers a new way to studying dynamics of translocations induced by gene editing at various endogenous genomic locations. The possibility of combining CRISPR LiveFISH with gene editing and other manipulation techniques (e.g., CRISPRi/a, epigenetic modification, CRISPR-GO) should further expand our capability to study the spatiotemporal dynamics of genome organization and rearrangements (16,20,43–47).

## Acknowledgements

The authors thank Stanford Cytogenetics Department for karyotyping, the Cell Sciences Imaging Facility for microscope usage, CHEM-H Macromolecular Structure Knowledge Center for protein purification facilities, and Coriell Cell Repositories for providing cell resources. The authors acknowledge support from Pew Scholar Foundation, Alfred P. Sloan Foundation, and Li Ka Shing Foundation. This work was supported from the National Institutes of Health Common Fund 4D Nucleome Program (Grant U01 EB021240).

## Author contributions

H.W. and L.S.Q. conceived of the idea. H.W. and L.S.Q. planned the experiments. H.W., M.N., D.Z., C.N., A.L. performed the experiments. H.W., C.N., T.D., and L.S.Q. analyzed the data. H.W., M.L. and L.S.Q wrote the manuscript. All authors read and commented on the manuscript.

## Materials and Methods

### Plasmid construction and protein purification

For expression of the dCas9-EGFP protein, dCas9-EGFP with two copies of nuclear localization signal (NLS) was subcloned from pHR-TRE3G-dCas9-GFP plasmid (16) into pET-based bacterial expression vector containing an N-terminal His-MBP-TEV tag (30). dCas9-EGFP was purified following established protocols for dCas9 proteins (30). dCas9 protein was obtained from Jennifer A. Doudna’s lab at UC Berkeley (36).

### Guide RNAs synthesis, purification and preparation

crRNAs fluorescently labeled at 5’ end were synthesized by IDT (Integrated DNA Technologies, Redwood City, CA). Unlabeled tracrRNAs were synthesized by IDT or through *in vitro* transcription (HiScribe™ T7 Quick High Yield RNA Synthesis Kit, NEB) using DNA templates containing a T7 promoter binding sequence and full length tracrRNAs, and purified with RNA Clean & Concentrator™ -100 (Zymo Research, Irvine, CA). We included a A-U flip in the original cr:tracrRNA backbone sequence based on previous reported improvement with the sgRNAs for imaging application (11,16). For editing gRNAs in translocation experiments, Alt-crRNAs and tracrRNAs were purchased from IDT.

Fluorescent crRNA and tracrRNAs at equal molar ratio were annealed in folding buffer (20 mM HEPES, pH 7.5 and 150 mM KCl), incubated at 95°C for 5 min, 70°C for 5 min, gradually cooled down to room temperature and supplemented with 1mM MgCl_2_, then kept at 40°C for 5 min and gradually cooled down.

### Cell culture

U2OS cells were cultured in DMEM with GlutaMAX (Life Technologies) in 10% Tet-system-approved FBS (Life Technologies). Patau Syndrome patient-derived amniotic fluid cells (ID: AG12070, trisomy 13) and normal amniotic fluid cells (ID: GM00957, apparently healthy) were obtained from the NIGMS Human Genetic Cell Repository at the Coriell Institute for Medical Research (Camden, NJ), cultured in AminoMax II (Thermo Fisher, 11269016) supplemented with 10% Tet-system-approved FBS.

Human primary T cells (peripheral Blood CD3+ Pan T Cells) were purchased from Stem Cell Express (Placerville, CA), activated using Dynabeads Human T-Activator CD3/CD28 beads (Gibco, Life Technologies), and cultured in X-VIVO 15 (Lonza Cat# BE02-053Q) with 10% AB Human Serum (Thermo Fisher) and IL-7, IL-15 and IL-2 (Peprotech).

All cells were cultured at 37°C and 5% CO_2_ in a humidified incubator.

### CRISPR imaging by plasmids delivery

U2OS and Patau Syndrome patient-derived amniotic fluid cells were transfected by pHR-TRE3G-dCas9-EGFP plasmid and pHR-U6-sgRNA (16). We used leaky expression of dCas9-EGFP under Doxycycline (Dox)-inducible TRE3G promoter to reduce background of dCas9-EGFP accumulation (16). Cells were plated one day ahead and cultured overnight. Before electroporation, cells were digested with trypsin and washed with DPBS (Thermo Fisher, 14200166). Electroporation was performed using Neon Transfection System 10µL kit (Thermo Fisher, MPK1025), in U2OS cells at 1400v/30ms/2pulses and in Patau Syndrome patient-derived amniotic fluid cells at 1400V/30ms/4pulses (48). Cells were imaged 16-24 h after transfection. Hoechst 33342 (Thermo Fisher, H3570) was added to cells at 0.1µg/ml to stain the nucleus before imaging.

### Live-cell delivery of fluorescent RNP complexes

RNP delivery by electroporation was performed using Neon Transfection System 10µL kit (Thermo Fisher, MPK1025) (48) and transfected cells were plated in 24 ibidi µ-plates for imaging. For T cell imaging, the plates were pretreated with 1mg/ml collagen (Corning Incorporated, 354236) for 1 h at 37°C to accelerate cell attachment. To increase transfection efficacy, U2OS and amniotic fluid cells were pre-treated with nocodazole (Sigma, M1404, final concentration: 40-100ng/ml) for 16 hours before transfection (25).

For RNP delivery of fluorescently labeled cr:tracrRNA complexes, 10-25 pmol of each labeled crRNA:tracrRNA were mixed with equal molar amount of dCas9-EGFP or dCas9 in transfection buffer R or T and then incubated at room temperature for 10 min to allow for RNP assembly. The assembled RNP complexes were transfected into 1-2 × 10^5^ suspended cells using the standard protocol for the Neon Transfection System 10µL kit. Electroporation was performed in U2OS cells at 1400v/15ms/4pulses, in activated T cells at 1400V/10ms/3pulses, and in amniotic fluid cells at 1400V/30ms/4pulses. The transfected cells were immediately plated into 24-well ibidi µ-plates containing pre-warmed culture medium for imaging. Hoechst 33342 was added to cells at 0.1 µg/ml to stain the nucleus 4h after transfection or right before imaging.

### Generation and visualization of endogenous translocation events

For generation and visualization of endogenous translocation events, Cas9 RNP complexes were assembled by mixing 31 pmol of Cas9 (IDT, Cas9) with two labeling gRNAs (Atto488-gRNA targeting Chr3 and Atto565-gRNA targeting Chr13, 7.5 pmol each) to visualize dynamics of targeted Chr3 and Chr13 loci, and two unlabeled editing gRNAs (gPPP1R1 and gSPACA7, 7.5pmol each) in transfection buffer R to create double stranded breaks (DSBs) near the two labeled loci on Chr3 and Chr13. Editing gRNAs were assembled using IDT synthesized Alt-crRNAs annealed with IDT tracrRNAs following IDT-provided protocols. The assembled RNP complexes were transfected into 2 × 10^5^ suspended cells using the standard protocol for the Neon Transfection System 10µL kit. Electroporation was performed in U2OS cells at 1400v/15ms/4pulses.

### *In vitro* gRNA binding and RNase assay, gel electrophoresis and western blotting

To test the target DNA-dependent protection of gRNAs in RNase-rich environment, 1.1 µM Atto565-cr:tracrRNAs targeting 17nt of LacO region were incubated with 1 µM of purified dCas9s in dCas9 binding buffer (20 mM Tris·HCl (pH 7.9), 100 mM NaCl, 5% glycerol, 0.1 mM DTT, and 1 mM MgCl_2_) at room temperature for 10 min to form fluorescent ribonucleoprotein (fRNP) complex. Then, the fRNP complex was diluted by 5-fold (final concentration: 0.22 µM gRNA and 0.2 µM dCas9) and incubated with 2.5 µM of on-target DNA or non-target DNA at 37°C for 30 min. After this, RNase A (10mg/ml, Thermo Fisher Scientific, EN0531) was added at 1:2500 dilution into each sample. After RNase A addition, samples were incubated at 37°C, collected at different time points, and immediately frozen in liquid nitrogen. Collected samples were analyzed using native gel electrophoresis (NativePAGE 4-16% Bis-Tris Gel) or denature gel electrophoresis (SDS-PAGE, Novex 4-20% Tris-Glycine Mini Gels, Thermo Fisher Scientific). For SDS-PAGE, samples were mixed with Tris-Glycine SDS sample buffer and NUPAGE reducing agent and heated at 95°C for 2min before sample loading.

To investigate how on-target DNA binding and DNA mismatches affect gRNAs properties within CRISPR complexes, the fRNP complex (0.22 µM gRNA and 0.2 µM dCas9) was incubated with 2.5 µM of on-target DNA, non-target DNA or DNA containing 2, 4, 6, 13 or 15 mismatches at PAM-distal sites at 37°C for 30 min. All samples were run on NativePAGE 4-16% Bis-Tris Gel to separate dCas9 bound gRNAs and unbound gRNAs (Thermo Fisher Scientific). Samples containing only dCas9, only Atto565-cr:tracrRNA, or fRNP complex without DNA at same concentrations were also included in the same gel as controls.

After native gel electrophoresis, the Atto565-cr:tracrRNA fluorescence was detected under Typhoon FLA 9500 gel imager (GE healthcare) using the 532nm laser and a LPG filter (>575nm). The samples were subsequently transferred to iBlot 2 Nitrocellulose membrane using iBlot 2 Dry Blotting System (Thermo Fisher Scientific) for western blotting. Anti-Cas9 antibody (Diagenode, C15200203, 1:1000 dilution), and HRP-conjugated anti-mouse antibody (1:2000) was used to detected the bands of dCas9 proteins.

To detect if on-target DNA binding affects the brightness of Atto565-gRNA fluorescence, we aliquoted 2 µl of each sample and measured their fluorescence emission spectrum in a low volume 384 well plate (Corning, CLS4514) using a Synergy H1 microplate reader (Biotek Inc.) with 488nm excitation.

### Karyotyping

The karyotype analysis of normal and PS patient-derived amniotic fluid cells were performed by the Cytogenetics Department at Stanford Health Care using standard methods. Cells were harvested by standard cytogenetic methodologies of mitotic arrest, hypotonic shock and methanol-acetic acid fixation, and metaphase preparations were subsequently analyzed by the GTW chromosome banding methodology.

### PCR, cloning and sequencing for translocation segments

U2OS cells were electroporated with Cas9 RNP complexes containing the two imaging gRNAs and the two editing gRNAs as described previously. At different time points (27h, 56h and 90h) after transfection, genomic DNA was extracted using DNAeasy Blood and tissue kit (Qiagen). To tested if translocation created chimeric chromosomes containing both CRISPR LiveFISH labeled Chr3 and Chr13 loci. 81 different pairs of primers were used to amplify translocation segments of different combinations. PCR reactions were performed using Kapa Hotstart PCR kit (Fisher Scientific) on BioRad T100 Thermal cycler under the following conditions: 95°C for 3 min; 50 cycles of 98°C 20s, 65°C 20s, 72°C 1 min, and final extension 72°C for 10 min. Among those primers, 4 pairs of them produced clean amplicons of expected sizes in treated samples (**Supplementary Figure 5a**). The PCR products were gel purified using Nucleospin Gel and PCR clean up kit, cloned into TOPO vectors using Zero Blunt TOPO PCR cloning kit (Thermo Fisher), and transfected into Stellar Competent cells (Takara/Clontech). For each PCR amplicon, six colonies were picked for plasmids extraction and sequencing, and three sequencing reactions were performed for each colony to confirm sequencing results.

### Microscopy and Image processing

Microscopic imaging was performed on a Nikon TiE inverted microscope equipped with 100×PLAN APO oil objective (NA=1.49), the 60×PLAN APO oil objective (NA=1.40) or the 60×PLAN APO IR water objective (NA=1.27), an Andor iXon Ultra-897 EM-CCD camera and 405nm, 488nm, 561nm and 642nm lasers. Images were taken using NIS Elements version 4.60 software by time-lapse microscopy with Z stacks at 0.2μm, 0,3 μm or 0.4μm steps. For live cell imaging, cells were kept at 37°C and 5% CO_2_ in a humidified chamber.

For long-term live cell imaging, microscopy was performed in Nikon TiE inverted microscope equipped with the 60×PLAN APO oil objective (NA=1.40). Images were taken every 20-30 minutes for 10-13 hours. To visualize the nucleus, transfected cells were stained by SiR-DNA nuclear staining kit (Cytoskeleton. Inc) for 2 hours before imaging. Image processing was performed in Fiji (image J) (49) or MetaMorph (Molecular devices, CA). For Z-stack images, the average of maximum or average intensity of several Z planes are shown in figures. Some images were processed using the “smooth” function in Fiji to reduce noise for visualization purposes only. To compensate for bleaching effects over time, genomic labeling images are auto-contrasted using MetaMorph in supplementary **Movies 1-5**.

### Immunostaining

To detect the presence of DSBs in the translocation experiment, U2OS cells were transfected with editing RNPs and LiveFISH fRNPs and fixed in 4% paraformaldehyde (PFA) 10 hours after transfection. Chr3 loci were imaged by Atto488-crRNA, and Chr13 loci were imaged by Atto565-crRNA. Immunostaining was performed in fixed samples with Anti-53BP1(Novus Biological, Cat.# NB100-304SS), and donkey anti-rabbit Alex Fluor 647 secondary antibody (ThermoFisher, Cat.# A-31573). Hoechst 33342 (blue) labels the cell nucleus.

### Flow cytometry fluorescence assays

For quantification of gRNA degradation kinetics via RNP delivery in living cells, U2OS were electroporated with 10 pmol of Atto565 labeled cr: tracrRNA were mixed with equal molar amount of dCas9-EGFP. At different hours after electroporation, U2OS cells were dissociated using 0.25% Trypsin EDTA (Life Technologies) and analyzed by flow cytometry on CytoFlex S (Beckman Coulter Life Sciences) using 488nm and 561 lasers. At least 10,000 cells were analyzed for each sample. To quantify relative fluorescence over time, the fluorescence of untreated U2OS cells is set to 0, while the fluorescence of transfected cells at time 0 (right after transfection) is set to 1. Technical duplicates in 5 experimental replicates were analyzed.

### Calculation of signal to background ratio

In **Fig.1F**, linescans were performed using the “linescan” function in MetaMorph and plotted in Prism 8 (Graphpad.com). Relative fluorescence was calculated by setting the average background fluorescent intensity in the nucleus as 1. Signal to background (S/B) ratio was calculated by dividing peak fluorescent intensity of labeled chromosome loci by average background fluorescent intensity in the nucleus.

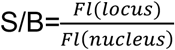

Where *Fl(Locus)* is the maximum fluorescent intensity at a labeled locus, and *Fl(nucleus)* is the average intensity within nucleus excluding the region of labeled loci.

### Dynamics tracking and 2D correlation analysis

Genomic loci tracking was performed using the TrackMate plugin (50) in Fiji. For tracking genomic loci, the estimated blob diameter was set between 0.5-1 μm, and for tracking nuclear movement, the estimated blob diameter was set between 10-15 μm. Linking max distance was set to 2 μm, and gap closing distance was set to 3 μm and gap closing max frame was set to 2. Tracks were analyzed in Excel and plotted with Datagraph or Excel. Mean square displacements (MSDs) were calculated using a mean square displacement analysis of particles trajectories (51) implemented in MATLAB (Mathworks Inc).

The 2D correlation between loci and nuclear movement was calculated using the following formula in Excel. Movement relative to the nucleus is calculated by subtracting x, y position of the nuclear center from the x, y positon of a given locus at each time point.

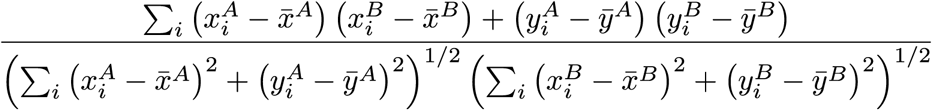

The formula calculates the 2D correlation between the dynamic tracks of loci A and B. where 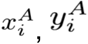 are the position of locus A at a given time point *i*,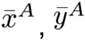, are the average position for locus A of all time points.

### Statistics Analysis

For **Fig. 1F**, *p* value was calculated using Wilcoxon two-sided *t*-test in Prism 7, and 47 loci in 17 cells were analyzed. For quantification of labeling efficacy (**Fig. 2D, E**, ≥100 cells were randomly selected from each of three independent experiments. For **Fig. 2D**, *p* value was calculated using a Student’s paired two-sided *t* test with unequal variance in Excel. Error bars show standard deviations in all figures.

## Data Availability

The authors declare that all data generated or analyzed during this study are either included in this published article (and its supplementary information files) or are available from the corresponding author on reasonable request.

## Supplementary Figures

**Supplementary Figure 1.**
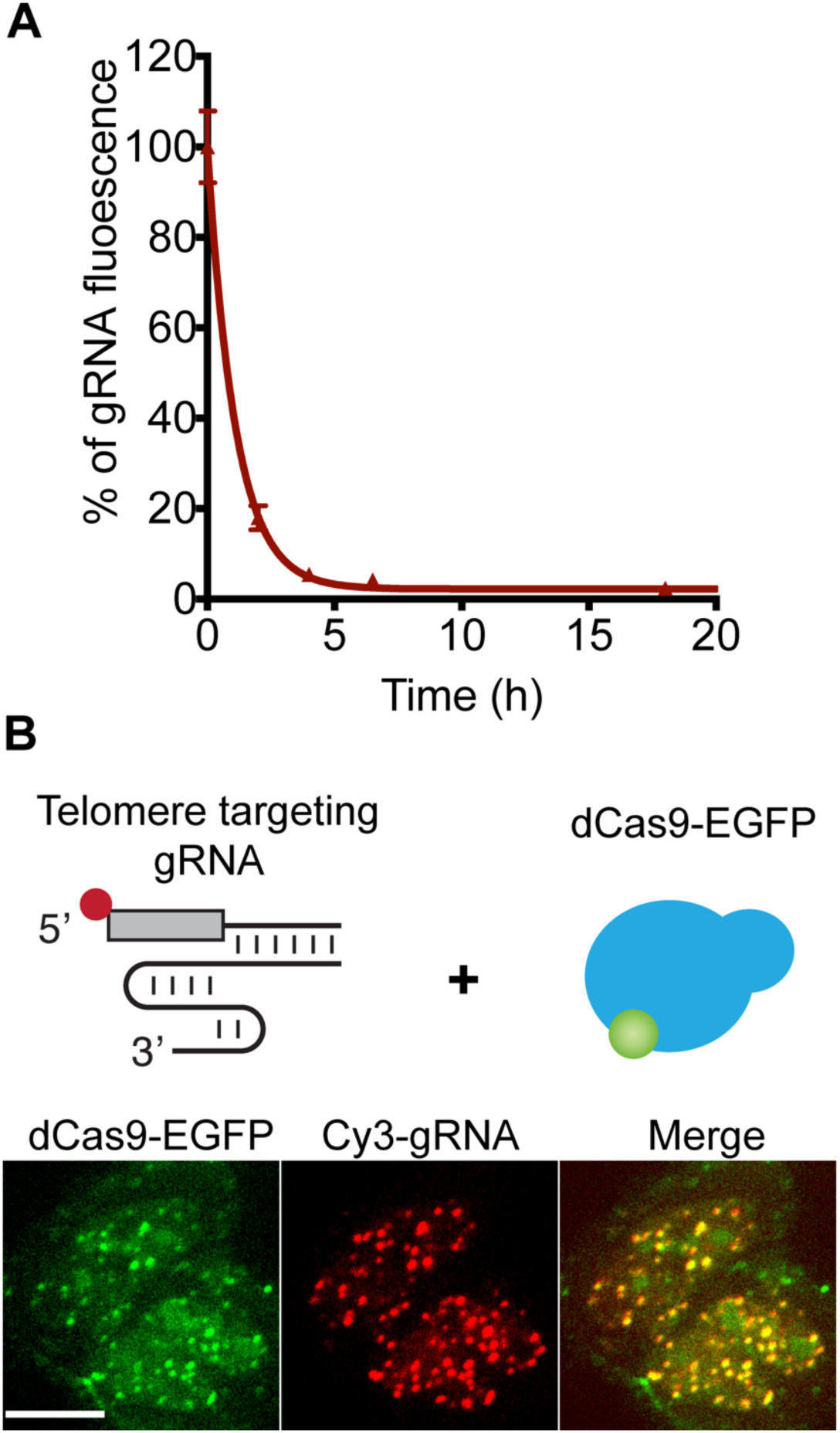
fRNP delivery of fluorescently labeled gRNAs and dCas9-EGFP in U2OS cells. (**A**) Flow cytometry analysis of Atto565-gRNA fluorescence after electroporation of Atto565-labeled gRNA and dCas9-GFP complex into U2OS cells. Time is shown as hours after electroporation. (**B**) Comparison of fluorescent gRNA (Cy3-gRNA) and dCas9-EGFP labeling for telomeres in U2OS cells using fRNPs. Both dCas9-EGFP (green) and telomere-targeting Cy3-gRNA (red) labeled telomeres, but there is less background in the Cy3-gRNA channel than in the dCas9-EGFP channel. Scale bar, 10 μm.

**Supplementary Figure 2.**
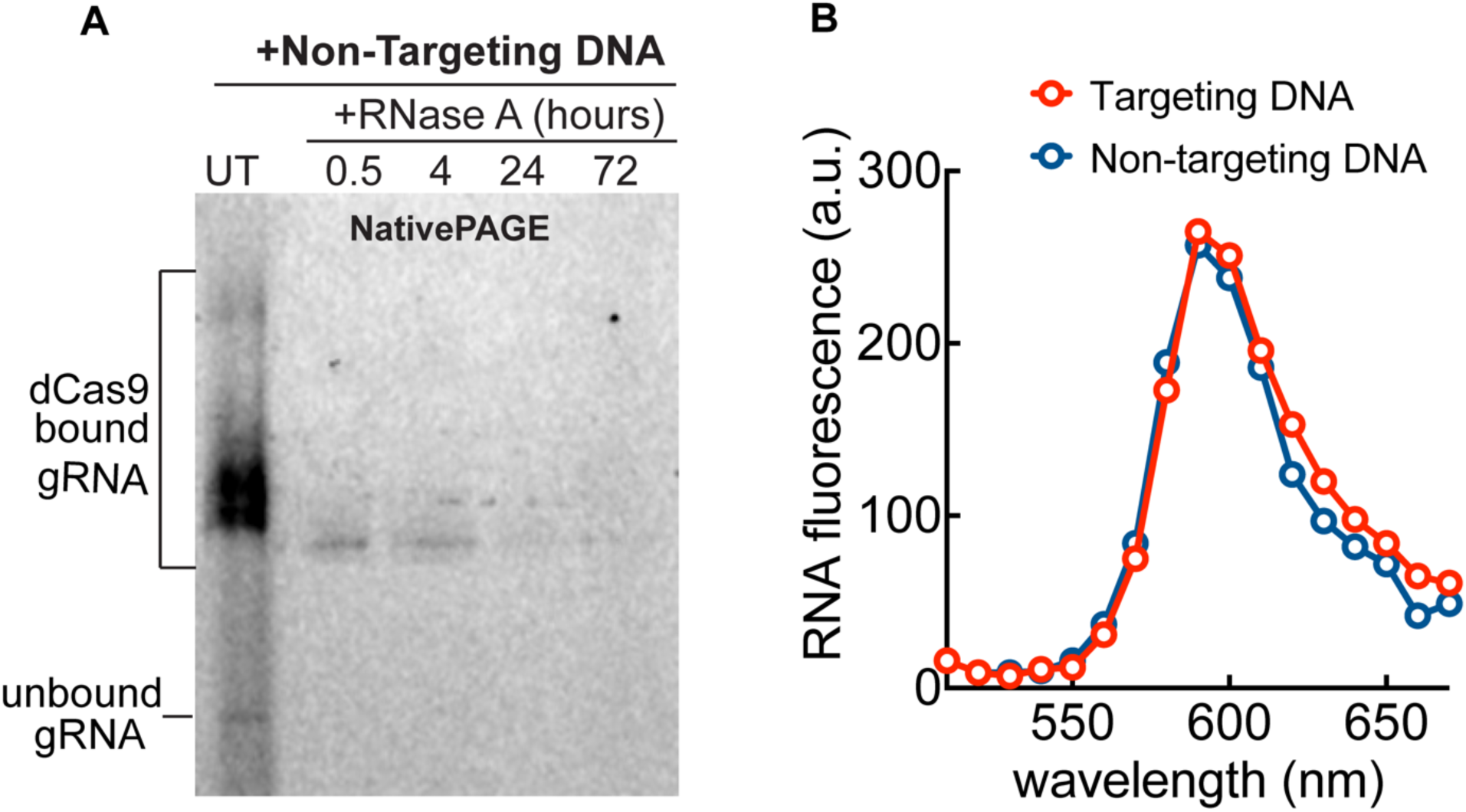
Biochemical assays studying gRNA properties in Cas9 RNP complex *in vitro*. (**A**) Native gel electrophoresis (NativePAGE) comparing fluorescent intensity of Atto565-gRNAs in complex with dCas9 and non-target DNA treated with RNase A for 0.5, 4, 24, and 72 hours (right four lanes). UT, Untreated with RNase. (**B**) Comparison of fluorescence emission spectrum of Atto565-gRNAs (gRNAs) incubated with dCas9 and on-target DNA (red) or non-target DNA (blue).

**Supplementary Figure 3.**
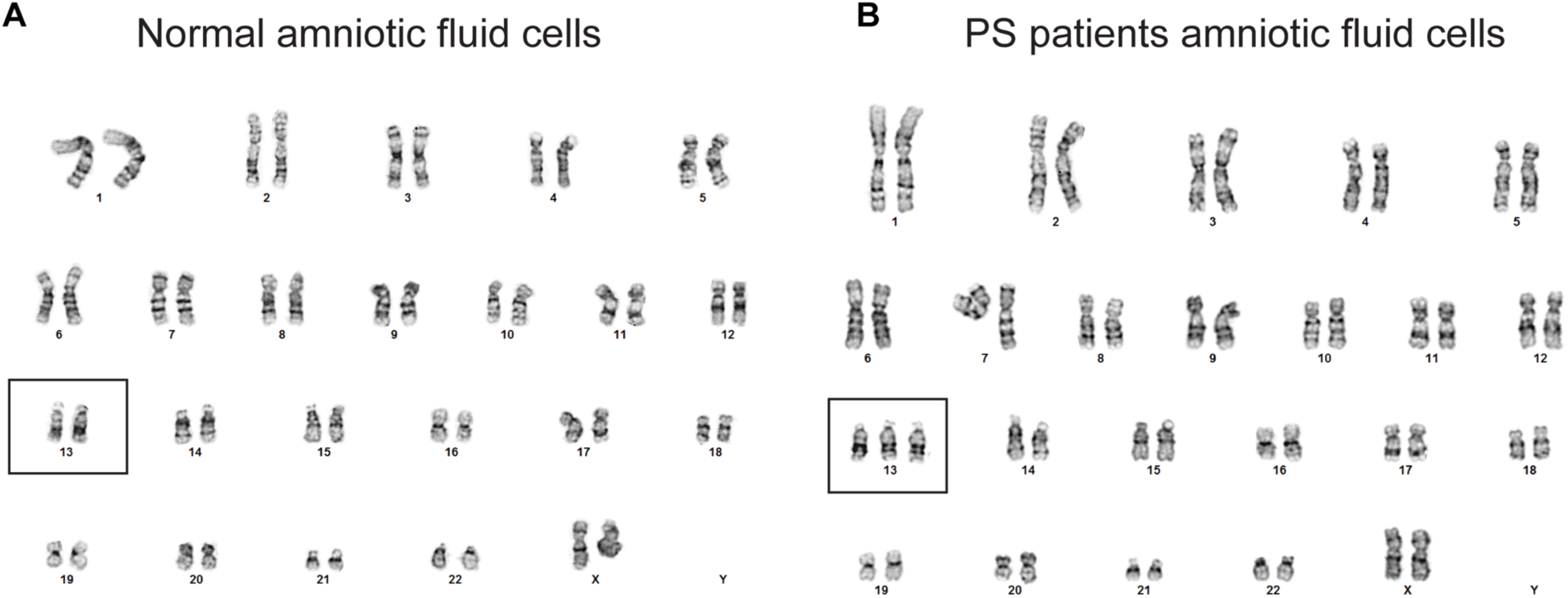
Controls for CRISPR LiveFISH. Karyotype analysis of normal (**A**) and PS patient-derived (**B**) amniotic fluid cells. Cells studied in **Figure 3B-C** were confirmed by karyotype analysis, showing 2 copies of Chr13 in normal amniotic fluid cells and 3 copies of Chr13 in PS patient-derived amniotic fluid cells.

**Supplementary Figure 4.**
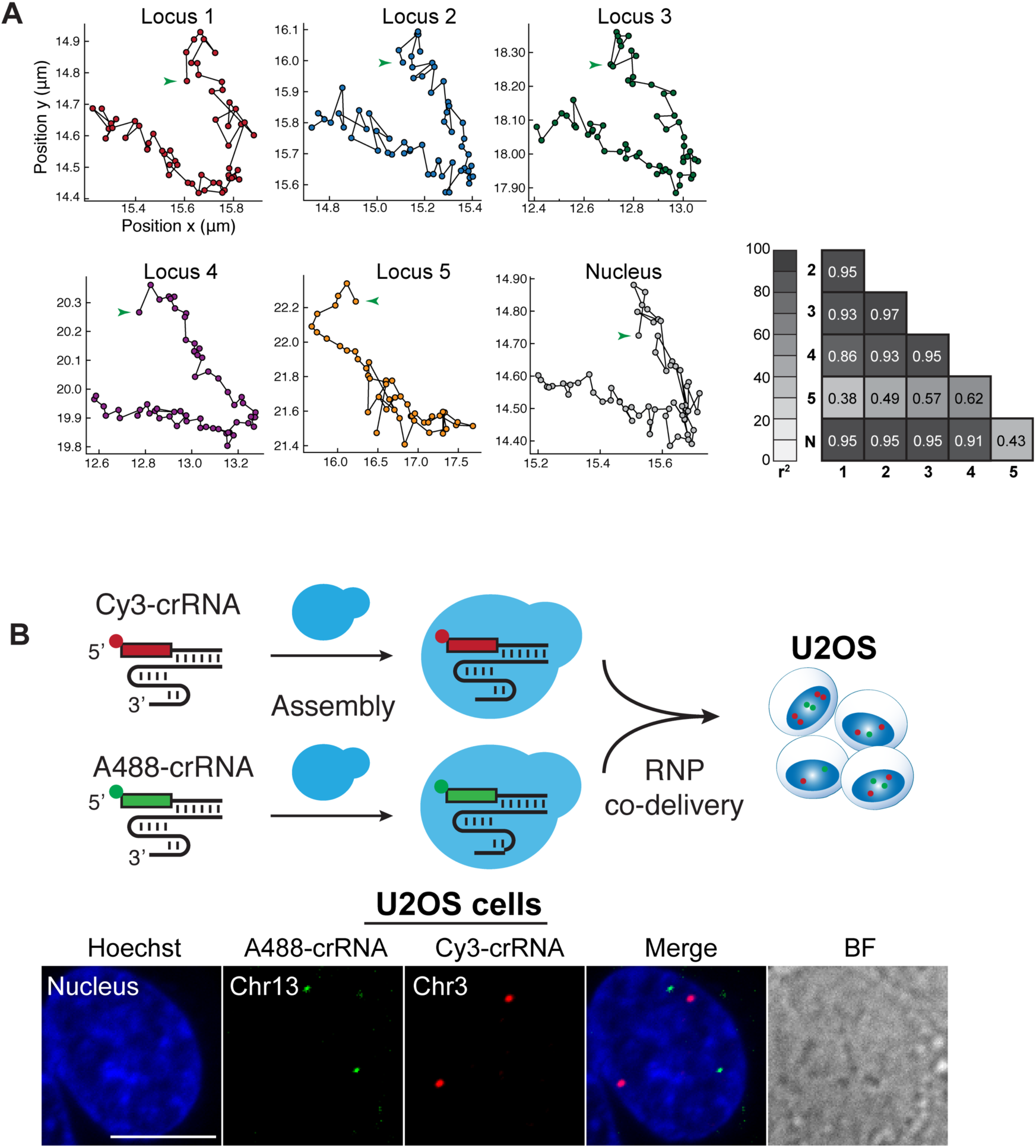
Dynamic analysis and multi-locus imaging of LiveFISH. **(A)** Dynamic trajectories of Chr13 loci in a normal amniotic fluid cell labeled with Atto565-gRNA (red). All five labeled loci and the nucleus movement are shown. Loci 1-4 are real genomic targets and locus 5 is cytoplasmic aggregation. The table shows relative 2-D correlation coefficient between each locus and with the nuclear movement. **Figure 3G** shows the mean square displacement (MSD) analysis. Scale bar, 10 μm. (**B**) CRISPR LiveFISH for multiple genomic loci imaging using differently labeled crRNAs in living U2OS cells. Simultaneous labeling of Chr13 (Atto488-crRNA, green) and Chr3 (Cy3-crRNA, red) by co-delivery of two CRISPR fRNP complexes in U2OS cells. Hoechst 33342 (blue) labels the cell nucleus. Scale bar, 10 μm.

**Supplementary Figure 5.**
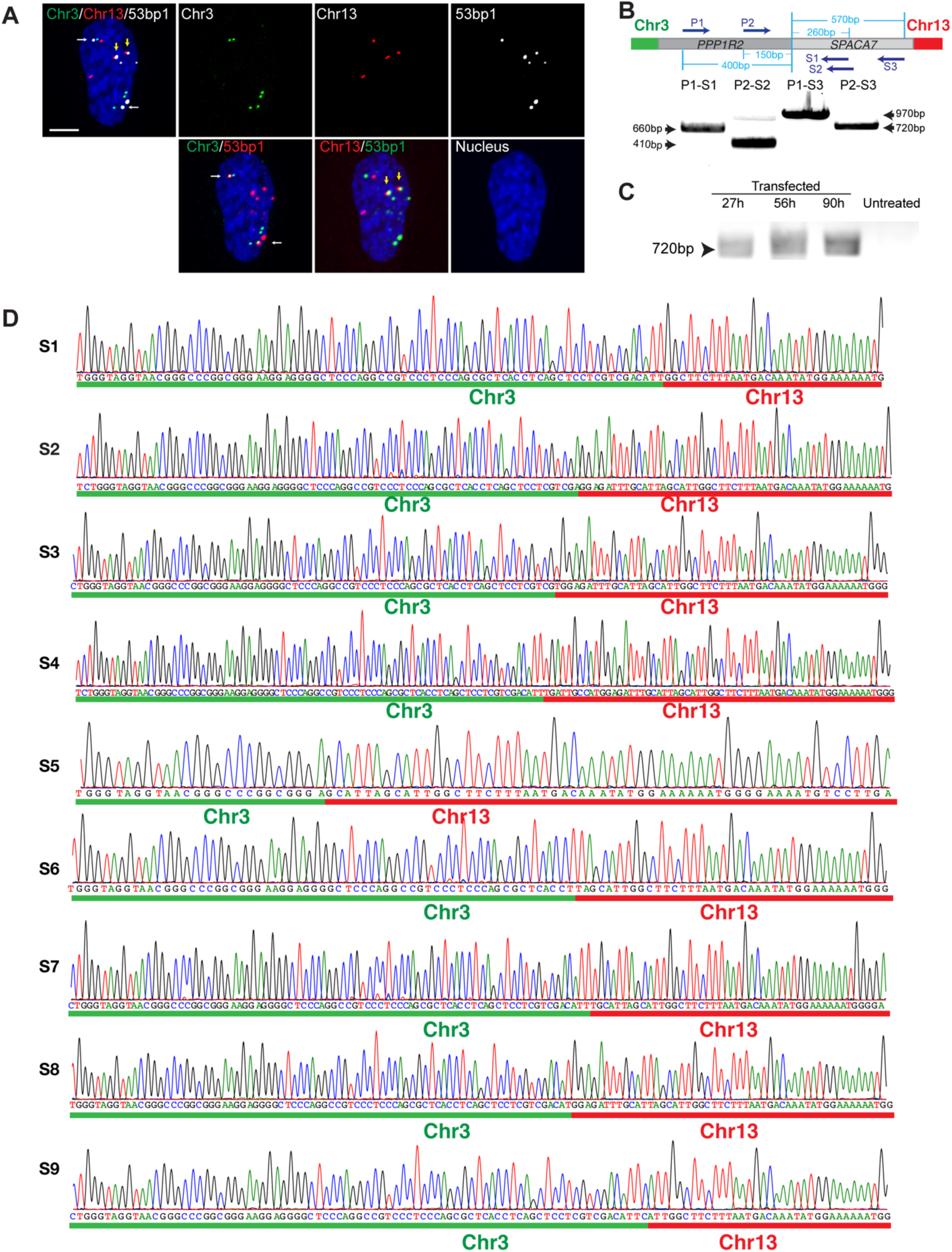
PCR and sequencing analysis confirming translocations between Chr3 and Chr13. **(A)** A representative U2OS cell showing co-localization of Chr13 and Chr3 loci with 53BP1 immunostaining. Scale bar, 10 μm.(**B**) DNA gel electrophoresis showing that multiple pairs of PCR primers produced translocated segments of expected sizes in treated cells. Top: The positions of the primer-binding sequences on the predicted translocated Chr3: Chr13 chromosome are shown: primers P1 and P2 targeting the *PPP1R2* gene on Chr3, and primers S1, S2 and S3 targeting the *SPACA7* gene on Chr13. Bottom: P1-S1, P2-S2, P1-S3 and P2-S3 show pairs of primers used to produce PCR product of expected sizes 660bp, 410bp, 970bp, and 720bp, respectively. (**C**) DNA gel electrophoresis compares PCR amplicons of translocated segments in RNP treated samples at different time points (27h, 56h and 90h post transfection) and untreated U2OS cells. (**D**) Sanger sequencing maps of translocation segments S1-S9 in **Figure 5G**. Green bars mark sequences mapped to Chr3 and red bars mark sequences mapped to Chr13.

**Supplementary Figure 6.**
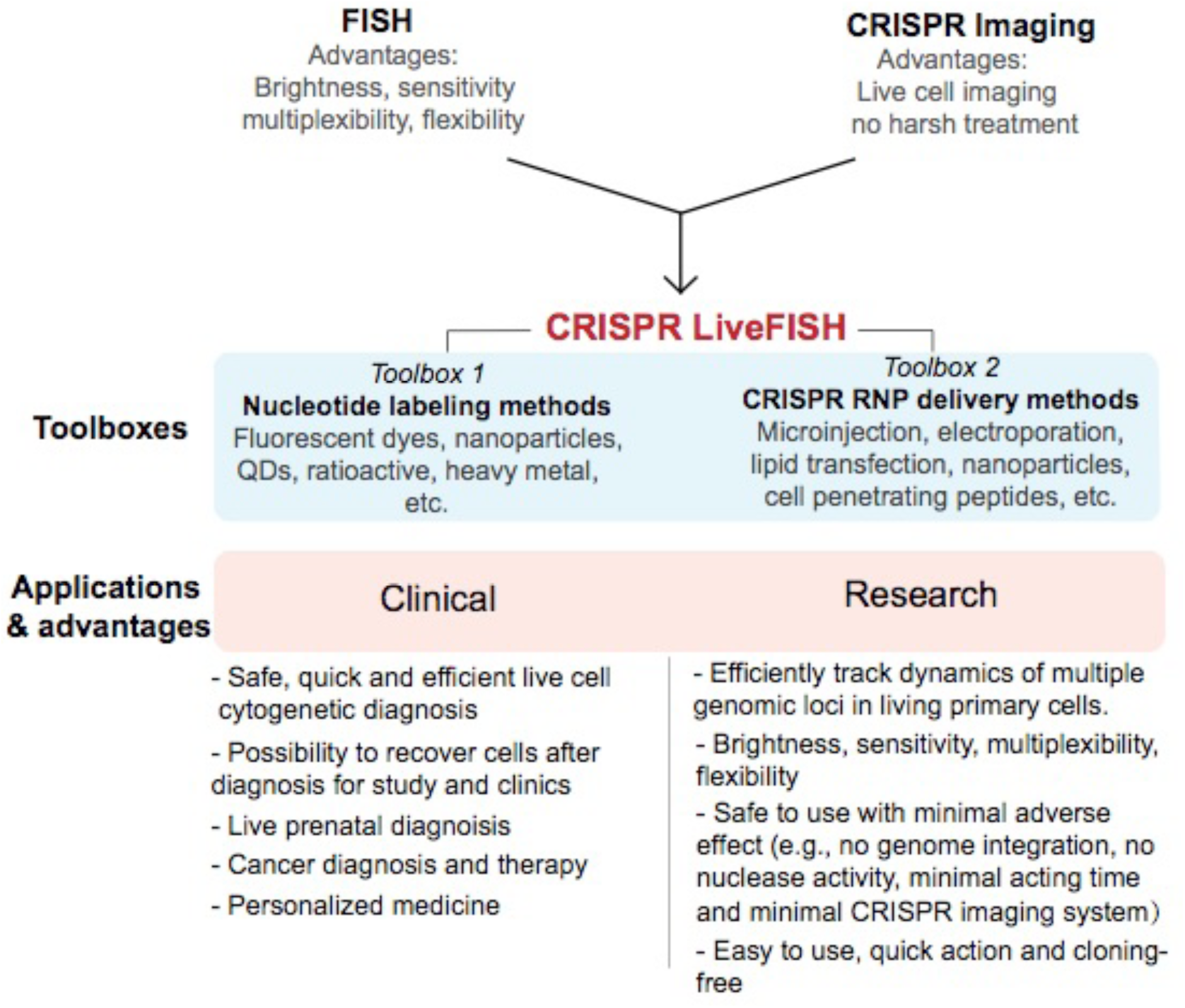
Advantages and potential applications of CRISPR LiveFISH. In CRISPR LiveFISH, the dCas9 protein can unwind DNA in living cells and specifically target fluorescently labeled RNA probes for genomic DNA hybridization and visualization. Thus, it combines the advantages of FISH (easy-to-use, high sensitivity, flexibility, multiplexibility) and CRISPR imaging (no harsh treatment, live-cell imaging). CRISPR LiveFISH provides a versatile platform that can be combined with diverse nucleotide labeling techniques and RNP delivery methods to optimize and expand its applications. Development of CRISPR LiveFISH may expand to the cytogenetic detection of various chromosomal abnormalities in cancer, infertility, and congenital disorders. CRISPR LiveFISH can also be used to study chromosome loci dynamics in living primary cells, which aids fundamental research to define the three-dimensional chromosomal organization in space and time during developmental and pathological processes.

**Supplementary Figure 7.**
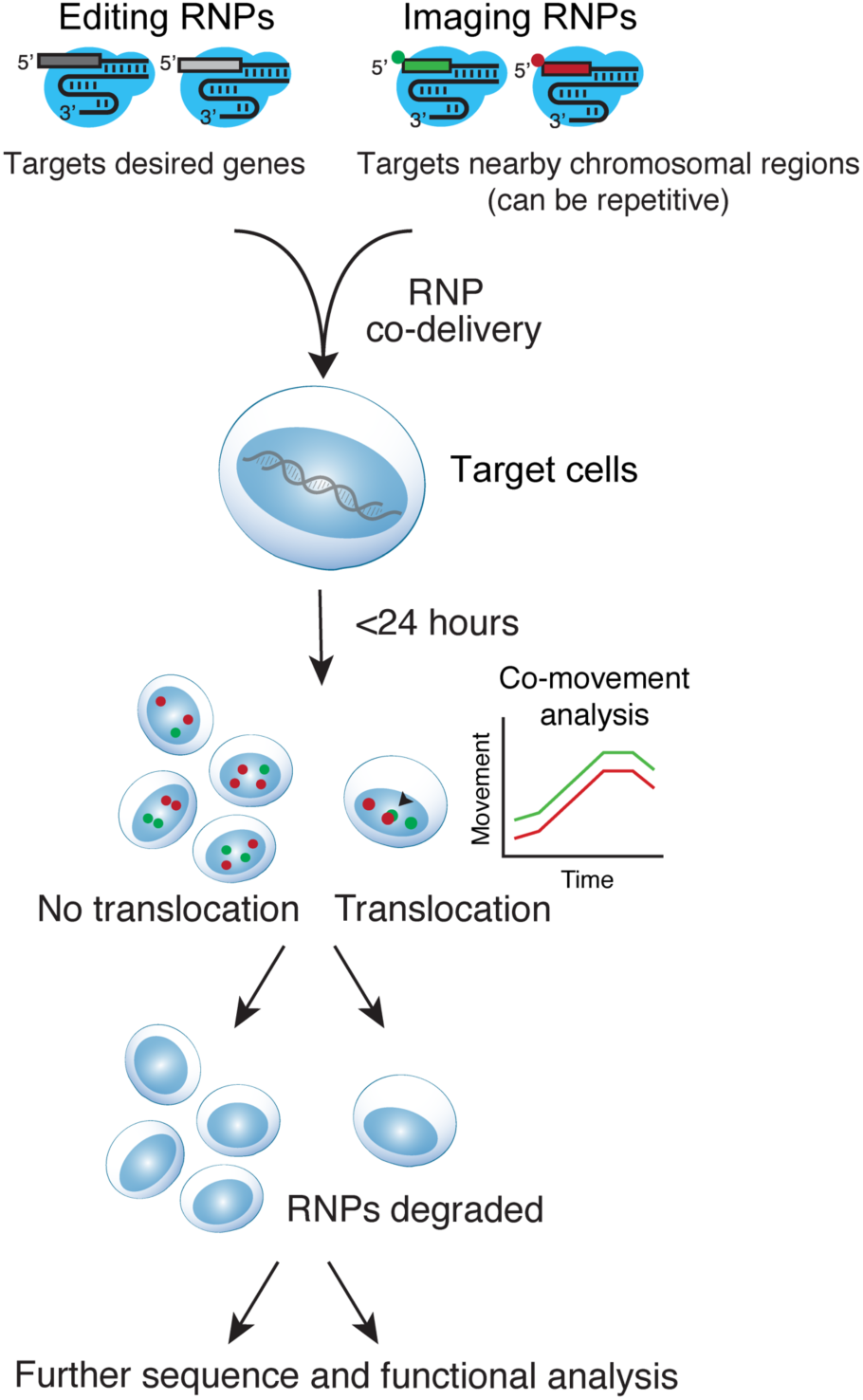
Real-time tracking of gene editing-induced chromosomal translocation. Visualization of endogenous chromosomal translocation induced gene editing via in a one-step reaction combining CRISPR LiveFISH and CRISPR editing. The imaging and editing RNP complexes are co-transfected into cells for editing and detection, and they are degraded after imaging, which minimizes adverse effect. The imaged cells remain alive, and thus can potentially be harvested for growth and further functional analysis.

## Supplemental Movies

**Movie S1. CRISPR LiveFISH tracks dynamics of Chr13 loci in a representative normal amniotic fluid cells**. Two Chr13 loci are labeled by Atto565-crRNA (gRNA, red). Hoechst 33342 (blue) labels the cell nucleus. Images were taken with seven Z confocal planes spaced by 0.2 μm every 18.5s, processed by projection of maximum intensity of all planes. Display rate 7 frames/s. See **Figure 2B**.

**Movie S2. CRISPR LiveFISH tracks dynamics of Chr13 loci in a representative PS patient-derived amniotic fluid cell**. Three Chr13 loci are labeled by Atto565-crRNA (gRNA, red). Hoechst 33342 (blue) labels the cell nucleus. Images were taken with seven Z confocal planes spaced by 0.4 μm every 17.1s. Display rate 7 frames/s. See **Figure 2C**.

**Movie S3. CRISPR LiveFISH enables multi-locus chromosome imaging in living primary human T lymphocytes**. Simultaneous labeling of Chr13 (Atto488-crRNA, green) and Chr3 (Cy3-crRNA, red) by co-delivery of two CRISPR RNP complexes in living primary human T cells. The recorded T lymphocyte moved slowly to the bottom right, with its movement being slowed down on a collagen pre-treated plate. Images were taken with 51 Z-confocal planes every 3 min (with a delay between frame 1&2), processed by projection of maximum intensity of all planes. Display rate 3 frames/s. See **Figure 2J** for another example with nucleus staining.

**Movie S4. Tracking dynamics of translocated endogenous chromosomes in living U2OS cells**. Simultaneous labeling of Chr3 (Atto488-gRNA, green) and Chr13 (Atto565-gRNA, red) by co-delivery of labeling gRNAs and cutting gRNAs in complex with Cas9 in living U2OS cells. A translocated chromosome is marked by a pair of closely located Chr3 and Chr13 sites in the center of nucleus. Images were taken with seven Z confocal planes spaced by 0.3 μm every 24 second, processed by projection of maximum intensity. Display rate 7 frames/s. See **Figure 3C**.

**Movie S5. Tracking formation of translocation between endogenous chromosomes in living U2OS cells**. Simultaneous labeling of Chr3 (Atto488-gRNA, green) and Chr13 (Atto565-gRNA, red) by co-delivery of labeling gRNAs and cutting gRNAs in complex with Cas9 in living U2OS cells. Images were taken with seven Z confocal planes spaced by 0.3 μm every 0.5 hours. Display rate 4 frames/s. See **Figure 3E**.

